# Postnatal maternal separation alters caregiving patterns but fails to promote anxiety- and depressive-like behaviour in C57BL/6N mouse dams

**DOI:** 10.64898/2026.09.22.753354

**Authors:** Victoria Mordvinova, Tamara Trowsse, Marie-Claude Audet

**Affiliations:** School of Nutrition Sciences, University of Ottawa, Ottawa, Canada; Department of Neuroscience, Carleton University, Ottawa, Canada; Department of Cellular and Molecular Medicine, University of Ottawa, Ottawa, Canada

**Author notes:** Correspondence to: Marie-Claude Audet, School of Nutrition Sciences, University of Ottawa, Ottawa, Ontario, K1H 8L1, Canada.

## Abstract

Stressors experienced during the postnatal period may increase vulnerability to mental health disorders, including postpartum depression and anxiety. In rodents, maternal separation (MS) has been shown to alter behaviours in dams, but these observations have been made almost exclusively in rats and thus robust mouse models of postpartum mental health disorders using this stressor remain limited. Here we first examined, in mice, the effects of MS on maternal caregiving and on anxiety- and depressive-like behaviours. As stress-induced inflammatory activation in the gut-brain axis has been associated with behavioural alterations in non-postpartum contexts, changes in pro-inflammatory cytokines and tight junction proteins in the brain and intestinal tract were also examined. Female C57BL/6N mice were assigned to a No Separation (NS) or a MS condition that consisted of 3-hour daily separation sessions from postnatal days (P) 2 to 14. Maternal care behaviours were assessed on P3 and P7, followed by anxiety- and depressive-like behaviours on P22 and P23, and the collection of the medial prefrontal cortex (mPFC) and colon on P24 to determine the expression of selected genes. Caregiving patterns in MS dams fluctuated throughout the early postnatal period. Although behaviours and gene expression outcomes remained unchanged by MS, grooming time in the splash test was correlated with the expression of different genes in the mPFC. These findings suggest that C57BL/6 mouse dams may be less sensitive to the actions of MS on behaviour and on brain and intestinal markers of inflammation and barrier permeability, at least when examined shortly after weaning.

## 1. Introduction

Postpartum depression and anxiety are prevalent mental health conditions in the Canadian population, with 23% of women reporting symptoms consistent with either of these disorders in the year after birth and 8% of women experiencing comorbid symptoms (Government of Canada, 2019). These conditions can disrupt the mother-infant relationship, with harmful consequences for both the mother and the child (Fairbrother et al., 2025; Somerville et al., 2015; Stewart & Vigod, 2019), highlighting the importance to advance our understanding of the processes that may contribute to their pathophysiology. The postpartum period is characterized by extensive physiological and behavioural adaptations that support the caregiving role and child development (Kim, 2016; Levin & Ein-Dor, 2023; Rincón-Cortés & Grace, 2020). It is thus not surprising that stressors experienced during this period are consistently reported as a major risk factor in both postpartum depression and anxiety development (Reid & Taylor, 2015; Yim et al., 2015), potentially because of their actions on hormonal and/or hypothalamic-pituitary-adrenal axis functioning (Hillerer et al., 2012; Levin & Ein-Dor, 2023; Schiller et al., 2015).

In rodent models, studies using maternal separation (MS), which can act both as an early life stressor for the offspring and as a postnatal stressor for the dam (Plotsky & Meaney, 1993), reported alterations in maternal care behaviours, such as arched-back nursing, grooming, licking, and pup retrieval (Baracz et al., 2020; de Almeida Magalhães et al., 2018; Rombaut et al., 2023) as well as increases in depressive- and anxiety-like behaviours in stressed dams (Boccia et al., 2007; Bousalham et al., 2013; Maniam & Morris et al., 2010). Notably, most of these studies have been conducted in rats and thus whether mouse behaviours are similarly affected by MS is incompletely understood. Additionally, little is known about the biological processes associated with postpartum behavioural phenotypes resulting from MS. In women, higher levels of pro-inflammatory cytokines in the plasma and cerebrospinal fluid have been associated with more severe symptoms of postpartum depression (Achtyes et al., 2020; Boufidou et al., 2009; Sha et al., 2022), suggesting that disturbances in inflammatory processes in circulation and the central nervous system could influence postpartum mental health. Women with inflammatory bowel disease were also reported to be at a higher risk of a new-onset mental disorder during the postpartum period (Vigod et al., 2019), raising the possibility that intestinal inflammatory disturbances could contribute to postpartum mental health disorders. Supporting this, perturbations to the signalling of inflammatory molecules between the intestinal environment and the brain are increasingly recognized as an important factor in the pathogenesis of mental health disorders (Audet, 2021; Cryan et al., 2019). In non-postpartum contexts, chronic stressors in mice increased barrier permeability and pro-inflammatory factors in selected intestinal and brain regions, contributing to behavioural impairments (Doney et al., 2023; Ménard et al., 2017; Russo et al., 2023; Szyszkowicz et al., 2017). However, whether these processes are disrupted by stressors in the context of postpartum mental health disturbances remains understudied.

Using C57BL/6N mice, we examined the effects of repeated postnatal stress in the form of MS on maternal caregiving during the early postpartum period and on anxiety- and depressive-like behaviours shortly after weaning. To determine whether behavioural alterations were accompanied by inflammatory and barrier permeability changes in intestinal and brain regions, the expression of selected pro-inflammatory cytokines and tight junction proteins in the medial prefrontal cortex (mPFC) and colon was quantified.

## 2. Methods

### 2.1 Animals

Eighteen primiparous female and nine male C57BL/6N mice, aged 6 to 8 weeks at their arrival, were used for breeding purposes (Charles River Laboratories, Montréal, Canada). All mice were housed in 27 cm x 21 cm x 14 cm polypropylene cages with a cardboard house, a cotton nestlet, and standard bedding, and maintained on a 12-h light-dark cycle, with the lights on from 0700 to 1900 h, at 23°C and 63% humidity, with food (Envigo, 2014 Teklad global 14% protein rodent maintenance diet) and water provided ad libitum. The University of Ottawa’s Animal Care Committee approved all experimental procedures (#4170), in compliance with the Canadian Council on Animal Care guidelines.

### 2.2 Summary of experimental procedures

Experimental procedures are summarized in Figure 1. Mice were housed individually upon their arrival and habituated to their new housing conditions for at least a week before mating using a harem breeding scheme. Upon the detection of a copulation plug, indicating Embryonic Day 0.5 (E0.5), female mice were returned to their home cages. Following the birth of the pups on postnatal day 0 (P0), litters were randomly assigned to an MS (*n* = 9) or a No Separation (NS; *n* = 9) condition. Litters were composed of equal numbers of female and male pups, ranging from 4 to 10 pups. MS litters experienced the stressor from P2 to P14 while NS litters were left undisturbed, after which all litters remained in standard housing conditions until weaning on P21. Body weight was recorded at different time points from pups’ birth to weaning. Caregiving behaviours were assessed on P3 and P7 whereas anxiety- and depressive-like behaviours were examined on P22 and P23. Mice were euthanized on P24, approximately twenty hours after the last behavioural test for the collection of the mPFC and colon.

**Figure 1:**
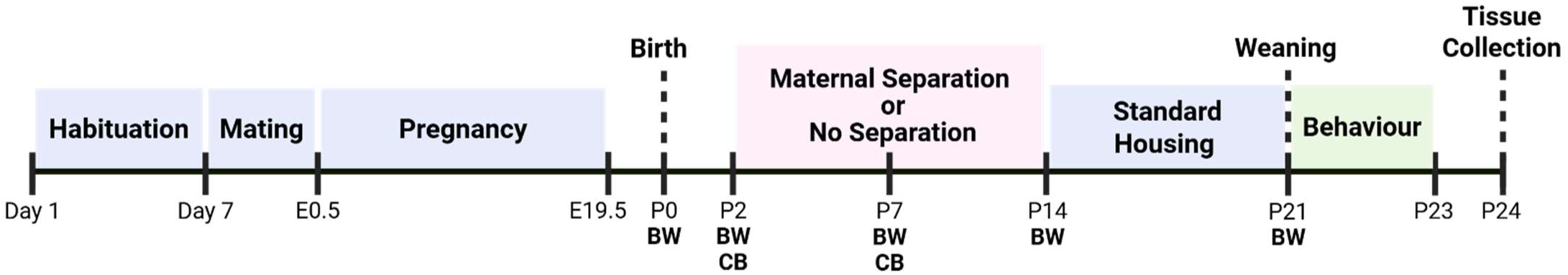
Timeline depicting the sequence of experimental manipulations from mice arrival on Day 1 to post-behavioural sample collection on postnatal day 24. BW: body weight; CB: caregiving behaviour; E: embryonic day; P: postnatal day.

### 2.3 Maternal separation procedures

Half of the litters underwent MS from P2 to P14, with the dams and their pups being separated for 3 hours each day (9:00 am to 12:00 pm), as described previously (Lippmann et al., 2007; Plotsky & Meaney, 1993). Briefly, pups were placed together in a new cage containing fresh bedding, a nestlet, and a cardboard house and kept in the same housing room, with a 30°C heating pad placed underneath the cage to reduce the risk of hypothermia. Dams were also placed in a new cage containing fresh bedding, a nestlet, a cardboard house, water, and chow, and transported to a different, empty housing room to prevent ultrasonic vocalisations between them and their pups. After each daily MS session, pups were reunited with their dam in their regular home cage until the next session. The other half of the litters (NS) remained in standard housing conditions and were not manipulated, except for routine weekly cage changes and body weight assessments.

### 2.4 Body weight measurements

The body weight of the dams was assessed at birth (P0), before the stressor (P2), mid-stressor (P7), at the end of the stressor (P14), and at weaning (P21) to examine the weight trajectory throughout different phases of the postpartum period. The percentage of change in weight from P2 to P14 was also calculated by dividing the difference between the two values by the body weight at P2 and multiplying by 100 to determine if the weight gained during the MS procedures differ between conditions.

### 2.5 Behavioural testing

Maternal care behaviours were assessed on P3 and P7. The elevated plus maze and open field tests were administered on P22 to assess anxiety-like behaviours, while the splash and tail suspension tests were performed on P23 to assess depressive-like behaviours, following an order from the least to most stressful. Mice were brought to the testing rooms approximately one hour prior to behavioural testing to habituate to their new environment. Each apparatus was cleaned with a Quato solution between trials.

#### 2.5.1 Maternal caregiving

Maternal care behaviours were assessed approximately 2 hours after the termination of the MS stressor or at equivalent times for NS litters. Briefly, dams were placed in a new cage in a room different from the one where testing was conducted, while their pups remained in their home cage positioned in a soundproof box. After 5 minutes, dams were reunited with their pups in their home cage, and their behaviours were recorded for another 5 minutes. The latency to first sniff pups as well as the time spent sniffing pups, moving pups and the nesting material, and in passive contact with pups (e.g., physical contact without direct interaction, for example sitting overtop of pups) were manually scored by a trained observer unaware of the experimental conditions. A score of total maternal care was also calculated by combining the time spent sniffing pups and moving pups and nesting material. A second observer, also blinded to the experimental conditions, scored 25% of the videos to determine interrater reliability.

#### 2.5.2 Elevated plus maze

The elevated plus maze was completed on P22 to assess the fear of open and elevated spaces, suggestive of anxiety-like behaviours (Komada et al., 2008; Walf & Frye, 2007). Briefly, mice were placed facing the intersection of open (6 cm × 75 cm) and closed (6 cm × 75 cm × 20 cm) perpendicular arms of a black opaque acrylic plastic apparatus raised 74 cm above the floor and allowed to explore the space for 5 minutes. The test was conducted under room lighting set at approximately 300 lux and a video-tracking software (EthoVision XT, Noldus) was used to determine the time spent in the open and closed arms and the total distance travelled.

#### 2.5.3 Open field test

The open field test was also conducted on P22, at least two hours after the elevated plus maze, to examine fearful behaviours in an open and lit space, suggestive of anxiety-like behaviours (Seibenhener & Wooten, 2015). Briefly, mice were placed in the bottom-right corner of a 45 cm x 45 cm x 45 cm opaque white acrylic plastic apparatus and allowed to explore for 10 minutes. The test was completed under approximately 300 lux of light and a video tracking software (EthoVision XT, Noldus) was used to determine the time spent in the center (15 cm x 15 cm) and the corners (10 cm x 10 cm) of the open field, as well as the total distance travelled.

#### 2.5.4 Splash test

The splash test was conducted on P23 to examine grooming behaviours (Isingrini et al., 2010). Reduced time spent grooming is reflective of a decreased motivation for self-care behaviours, as seen in human depression (Cathomas et al., 2015), whereas increased grooming time is indicative of hyperarousal (Kalueff et al., 2016). Mice were sprayed twice on their dorsal coat with a 10% sucrose solution, placed in an empty housing cage, and grooming behaviours were videotaped for 10 minutes. The test was conducted in a room with lighting at approximately 530 lux. Videos were scored manually by a trained observer unaware of experimental conditions to determine the total time spent grooming and the total number of grooming sessions. Based on the visual detection of subtle grooming differences in MS dams during manual scoring, the time spent in grooming sessions lasting over 3 seconds, termed sustained, was also determined. A second observer blinded to the experimental conditions scored 25% of videos to assess interrater reliability.

#### 2.5.5 Tail suspension test

The tail suspension test was performed last on P23, at least two hours after the splash test, to examine how long a mouse remains immobile in an inescapable situation. Increased immobility time is thought to be an indicator of passive coping responses to stress, a symptom found in human depression (Cryan et al., 2005; Steru et al., 1985). The tail suspension test was conducted using an apparatus that included an interface cabinet, a user interface software, and a load-cell amplifier hardware (Tail Suspension Test Cubicle [SOF-821], Med Associates Inc) under approximately 100 lux of light. Mice were attached to an aluminum bar at the end of their tail with surgical tape to prevent them from escaping the suspended position for 6 minutes. The aluminum bar was attached to a strain gauge and data were recorded by software (Tail Suspension SOF-821), allowing the determination of immobility time.

### 2.6 Brain and intestinal tissue collection

On P24, approximately twenty hours after the tail suspension test, mice were euthanized using rapid decapitation. Immediately afterward, whole brains were removed, placed on a cold stainless-steel matrix (2.5 cm x 3.75 cm x 2.0 cm, with slots spaced at approximately 500 μm) positioned on an ice plate, and sectioned coronally to collect the mPFC. In parallel, the gastrointestinal tract was removed from the abdominal cavity, placed on a nuclease-free surface on top of an ice plate, and the whole colon was dissected and emptied. All samples were placed in nuclease-free tubes positioned on dry ice immediately after their collection, and subsequently stored at −80°C.

### 2.7 Reverse transcription-quantitative polymerase chain reaction (RT-qPCR)

Total RNA was extracted from homogenized mPFC and colon samples using the manufacturer’s instructions (EZ-10 Spin Column Total RNA Miniprep Kit, Bio Basic Inc). A representative subset of mPFC and colon samples from each group was assessed for RNA quality using the Agilent Fragment Analyzer (Stem Core, University of Ottawa), with all tested samples demonstrating high RNA quality (average RNA integrity number score of 9.8). Concentration and purity were then assessed using a NanoDrop^TM^ One Spectrophotometer (Thermo Fisher Scientific) and only RNA samples with 260/280 ratios between 1.80 and 2.20 were included in subsequent analyses. The total RNA was converted into complementary DNA (cDNA) using iScript^TM^ Reverse Transcription Supermix (BioRad) and a T100 Thermal Cycler (BioRad). cDNA samples were analyzed by RT-qPCR with SsoAdvanced Universal SYBR Green Supermix (BioRad) and a CFX96 Touch Real-Time PCR Detection System (BioRad). Samples were analyzed in triplicates, and glyceraldehyde 3-phosphate dehydrogenase (*Gapdh*) was used as a reference gene. Fold changes in the mRNA expression of genes of interest in the mPFC and colon were determined using the −2^ΔΔCt^ method (Livak & Schmittgen, 2001; Schmittgen & Livak, 2008) relative to the NS group. Primer sequences are included in Supplementary Table 1.

### 2.8 Statistical analyses

Statistical analyses were conducted in GraphPad Prism (version 10.6.0). All data were tested for normality using the Shapiro-Wilk test and homogeneity of variance using the Levene’s test. Potential outliers were assessed using the robust regression and outlier removal (ROUT) method (Q = 1%). A limited number of samples were excluded from analyses due to technical difficulties with the equipment used for behavioural testing or poor RNA concentrations and/or purity, and thus the sample size and degrees of freedom differ between outcomes. A repeated two-way analysis of variance (ANOVA) with Stressor (NS versus MS) as a between-group factor and Time (P0, P2, P7, P14, P21) as a within-group factor was used to compare body weights across postpartum time points. Follow-up analyses to detect within-group differences were conducted using Bonferroni’s tests. Differences between the NS and MS groups in all other outcomes were assessed using independent *t*-tests for normally distributed data, Mann-Whitney *U* tests for data that did not meet the assumption of normality, or Welch’s *t*-tests for data that did not have the same variance. Spearman correlation analyses were conducted to examine associations between behavioural and biological measures. The alpha level for significance was set at 0.05 for all analyses.

## 3. Results

### 3.1 Body weight across the postpartum period

As seen on Figure 2A, changes in body weight from parturition to weaning were comparable in NS and MS dams (Time: F_(3.08, 49.26)_ = 56.19, *p* < 0.0001; Stressor and Stressor x Time: *p*’s > 0.05). Follow-up comparisons confirmed that the body weight increased between P0 and P14 (*p* < 0.0001) and then dropped from P14 to P21 (*p* < 0.0001), irrespective of the stressor condition. The weight gained between P2 and P14, encompassing the duration of the stressor, did not differ between NS and MS dams (Figure 2B; *p* > 0.05).

**Figure 2:**
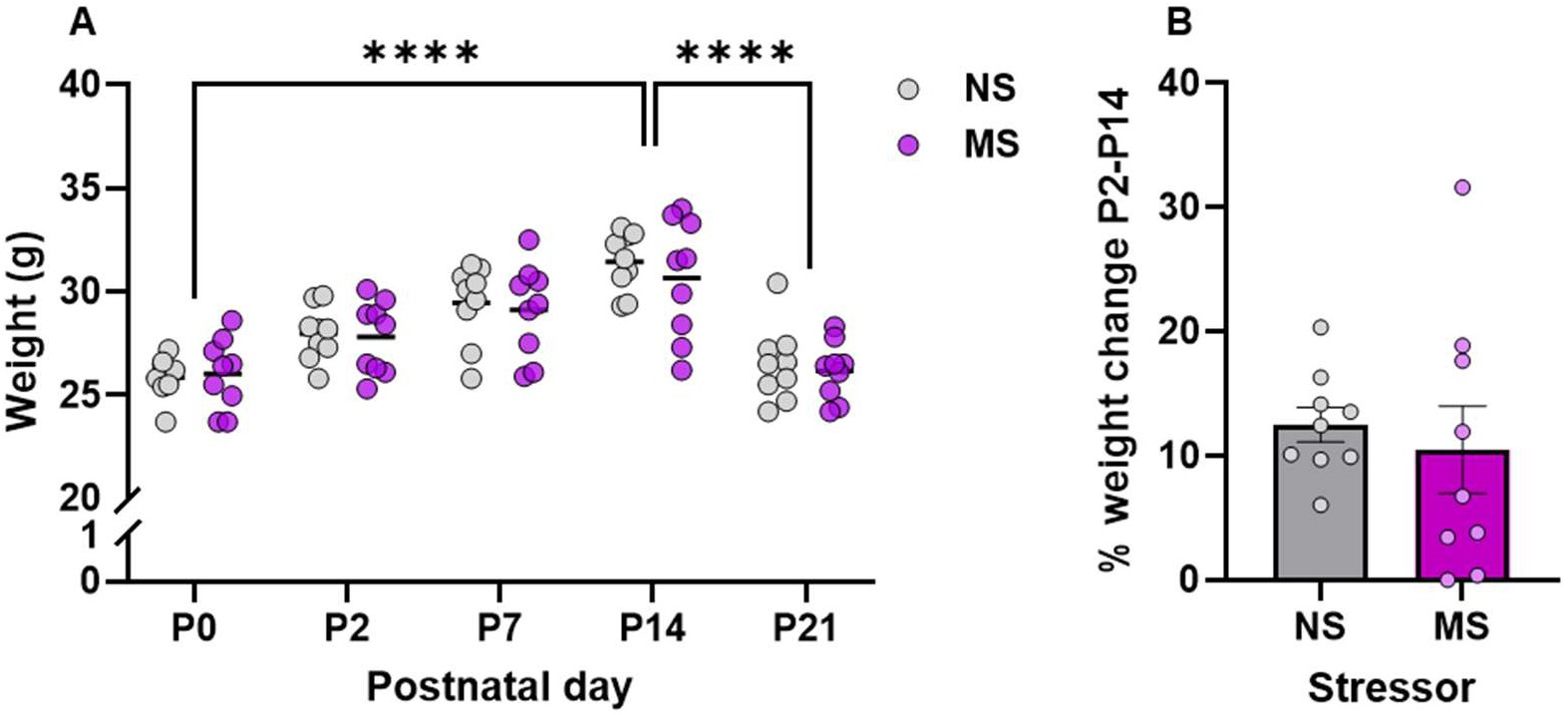
Body weight assessed in non-separated (NS) and maternally separated (MS) dams. Measures include body weight measurements at different time points during the postpartum period, including postnatal day 0 (P0), P2, P7, P14, and P21 (**A**) and the percentage in body weight change across the stressor period from P2 to P14 (**B**). Dots represent individual dams, and horizontal lines represent the median. NS (*n*= 9), MS (*n* = 9).

### 3.2 Maternal care behaviours during the early postpartum period

On P3, MS dams took more time to go sniff their pups after a 5-minute separation session (Figure 3A; *t*_(10)_ = 2.838, *p* = 0.018) and spent more time in passive contact with them during the 5-minute reunification that followed (Figure 3E; *t*_(10)_ = 2.245, *p* = 0.051), but these effects were not apparent anymore on P7 (Figure 3F,J; *p*’s > 0.05). In contrast, the lower time spent moving pups and the nesting material (Figure 3H; *U* = 9.000, *p* = 0.026) and the tendency to invest less time in maternal care overall (Figure 3I; *U* = 12.000, *p* = 0.072) in MS dams on P7 were not observed on P3 (Figures C,D; *p*’s > 0.05). Finally, MS dams did not differ from NS dams in terms of time spent sniffing pups on either P3 or P7 (Figures 3B,G; *p*’s > 0.05).

**Figure 3:**
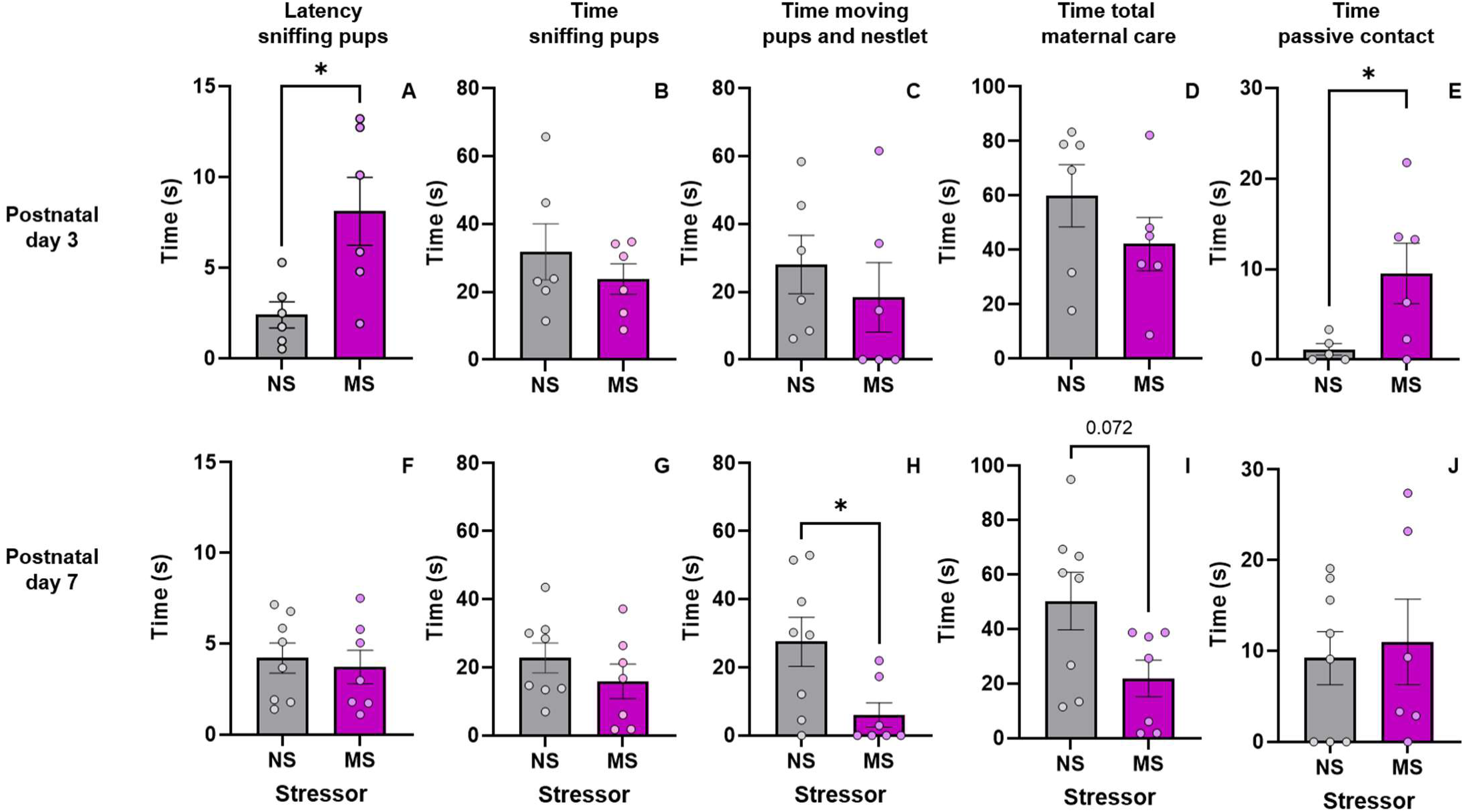
Maternal care behaviours assessed on postnatal day 3 (upper panels) and postnatal day 7 (lower panels) in non-separated (NS) and maternally separated (MS) dams upon reunification with their pups following a 5-minute separation session. Behaviours include the latency to first sniff their pups (A, F), the time spent sniffing their pups (B, G), the time spent moving their pups and the nesting material (C, H), as well as time engaged in maternal care behaviours (D, I) and in passive contact (E, J) with their pups. Dots represent individual dams, and bar plots and error bars represent means ± SEM. Postnatal day 3: NS (*n* = 6), MS (*n* = 6); Postnatal day 7: NS (*n* = 8), MS (*n* = 7). \**p* < 0.05 relative to NS.

### 3.3 Post-weaning anxiety-like and depressive-like behaviours

No differences were detected between NS and MS dams in the two tests used to assess anxiety-like behaviours and the two tests used to evaluate depressive-like behaviours. As shown in Figure 4A-C, the time spent in the open and closed arms as well as the total distance traveled in the elevated plus maze were similar across groups (*p*’s > 0.05). Likewise, the time spent in the center and in the corners in the open field test as well as the total distance travelled throughout the arena were unaffected by the MS stressor (Figure 4D-F; *p*’s > 0.05). In the splash test, the two groups did not differ in the total time spent grooming and the number of grooming bouts initiated (Figures 5A,C; *p*’s > 0.05). However, MS dams tended to spend more time in sustained grooming bouts, defined as grooming episodes lasting longer than 3 seconds, than NS dams (Figure 5B; *t*_(16)_ = 1.947, *p* = 0.069). Finally, the time spent immobile during the tail suspension test did not differ between NS and MS dams (Figure 5D; *p* > 0.05).

**Figure 4:**
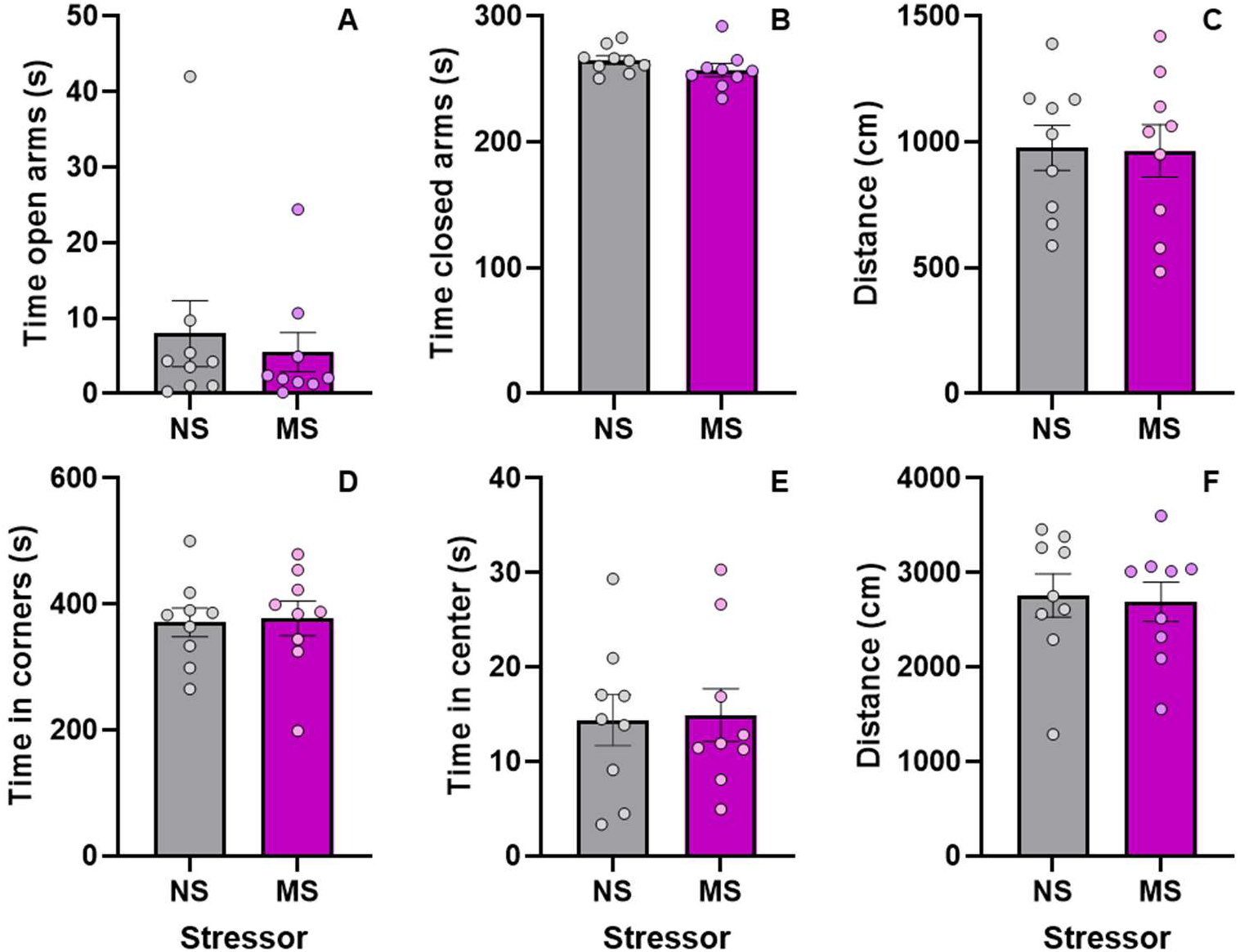
Anxiety-like behaviours were assessed on postnatal day 22. Time spent in the open (A) and closed (B) arms of the elevated plus maze and total distance travelled (C) in the apparatus as well as time spent in the center (D) and in the corners (E) of the open field test and total distance travelled (F) in the apparatus in non-separated (NS; *n* = 9) and maternally separated (MS; *n =* 9) dams. Dots represent individual dams, and bar plots and error bars represent means ± SEM.

**Figure 5:**
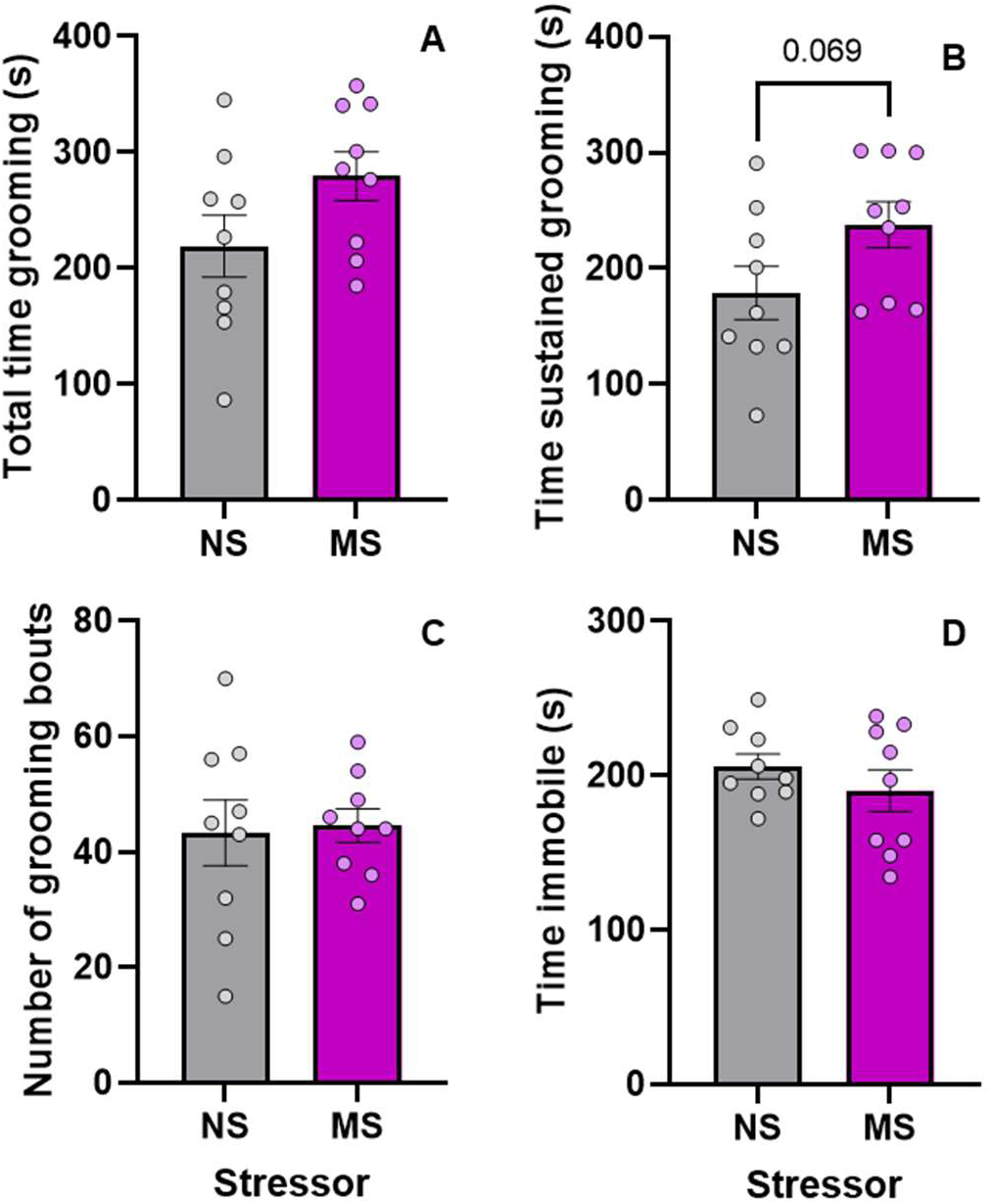
Depressive-like behaviours were assessed on postnatal day 23. Total time spent grooming (A), time engaged in grooming episodes longer than 3 seconds (termed sustained) (B), and frequency of grooming episodes (C) in the splash test as well as immobility time in the tail suspension test (D) in non-separated (NS; *n* = 9) and maternally separated (MS; *n* = 9) dams. Dots represent individual dams, and bar plots and error bars represent means ± SEM.

### 3.4 Gene expression of pro-inflammatory cytokines and tight junction proteins in the medial prefrontal cortex and colon

No differences were observed between the NS and MS groups in either the mRNA expression of pro-inflammatory cytokines or of tight junction proteins in the mPFC and colon (Table 1; *p*’s > 0.05). Because of the important dispersion of the behavioural and gene expression data within each of the NS and MS groups, we examined relationships between the main behavioural metrics determined during caregiving observation sessions and the elevated plus maze, open field, splash, and tail suspension tests and mPFC and colon pro-inflammatory cytokine and tight junction protein gene expression among the full sample of dams (including all mice from the NS and MS groups). As depicted in Figure 6, the time spent grooming was positively correlated with prefrontal TNF-α expression (*r* = 0.588 *p* = 0.038) but negatively correlated with IL-6 (*r* = −0.718, *p* = 0.016) and Ocln (*r* = −0.621, *p* = 0.027) in the same brain region. No other statistically significant correlations between behavioural and mRNA expression measures were identified (*p*’s > 0.05).

**Figure 6:**
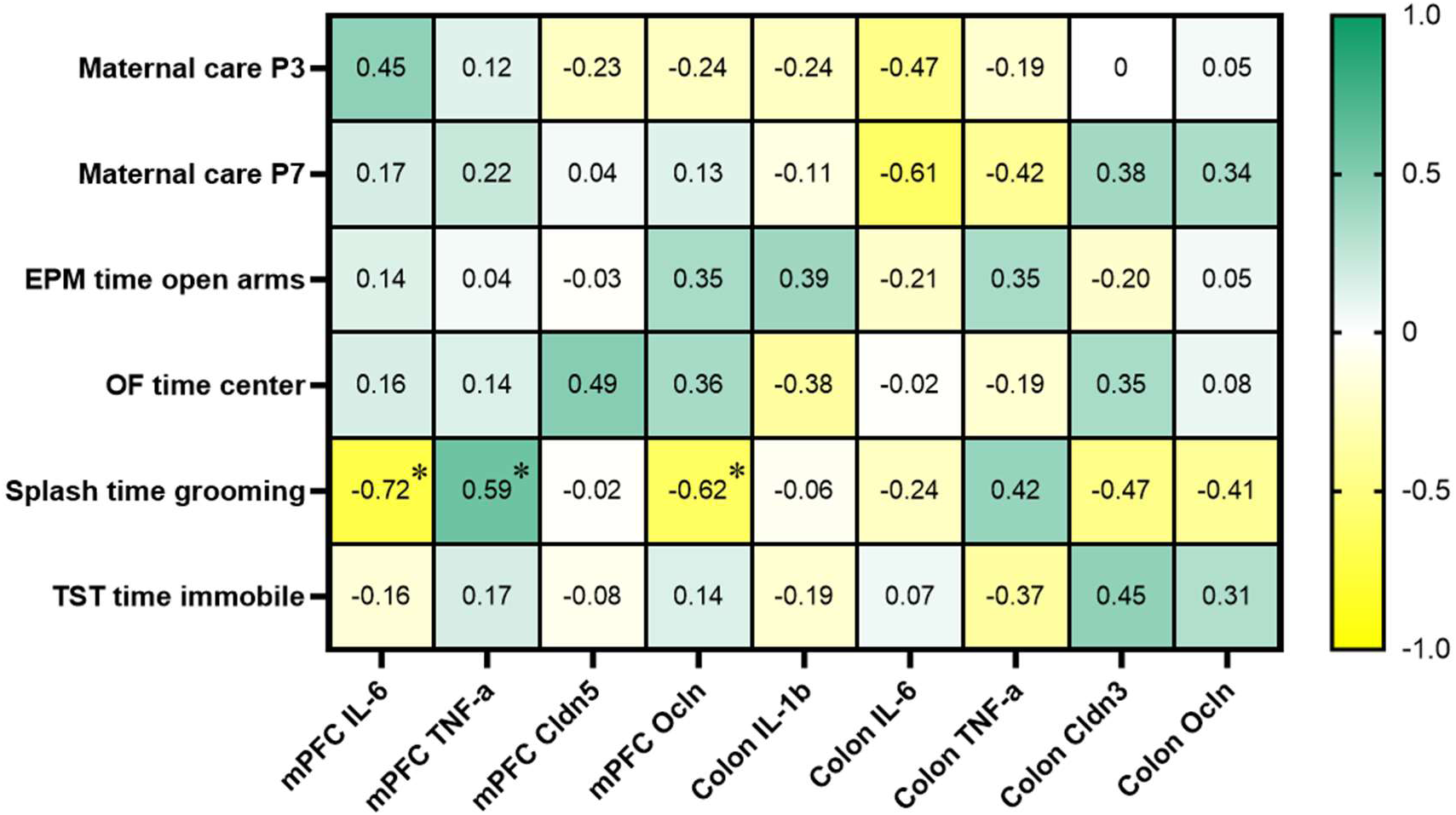
Spearman correlations coefficients between behaviours determined during caregiving observation sessions (total care), the elevated plus maze (EPM), open field (OF), splash, and tail suspension (TST) tests and fold changes in the mRNA expression for interleukin (IL)-6, tumour necrosis factor (TNF)-α, IL-1β, claudin 5 (Cldn5), Cldn3, and occludin (Ocln) in the medial prefrontal cortex (mPFC) and the colon. \**p* < 0.05.

**Table 1.**
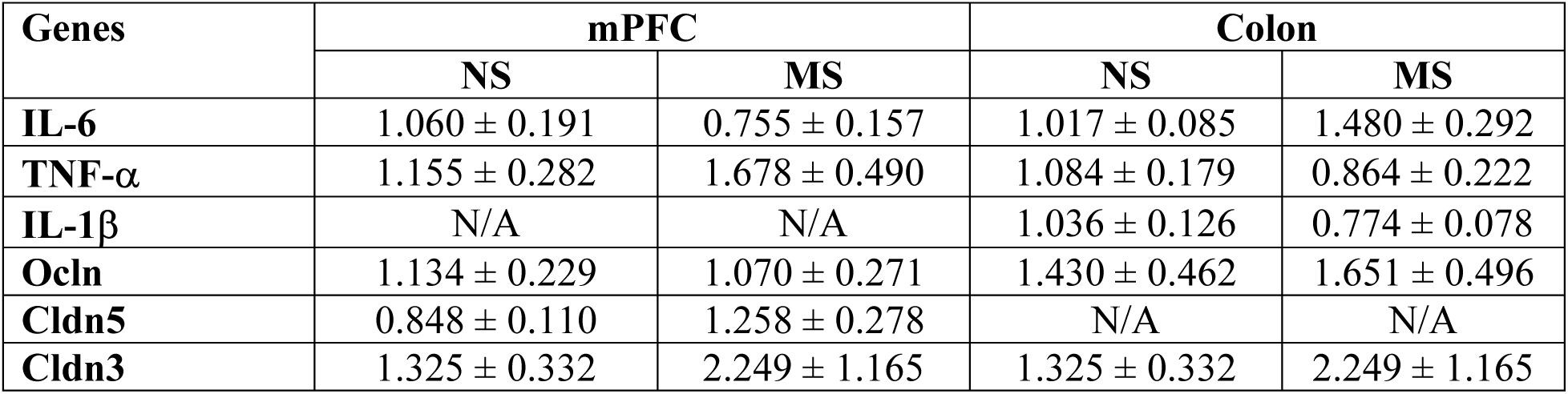
Gene expression of pro-inflammatory and tight junction markers in the mPFC and colon.

| **Genes** | **mPFC** |  | **Colon** |  |
| --- | --- | --- | --- | --- |
|  | **NS** | **MS** | **NS** | **MS** |
| **IL-6** | 1.060 ± 0.191 | 0.755 ± 0.157 | 1.017 ± 0.085 | 1.480 ± 0.292 |
| **TNF-α** | 1.155 ± 0.282 | 1.678 ± 0.490 | 1.084 ± 0.179 | 0.864 ± 0.222 |
| **IL-1β** | N/A | N/A | 1.036 ± 0.126 | 0.774 ± 0.078 |
| **Ocln** | 1.134 ± 0.229 | 1.070 ± 0.271 | 1.430 ± 0.462 | 1.651 ± 0.496 |
| **Cldn5** | 0.848 ± 0.110 | 1.258 ± 0.278 | N/A | N/A |
| **Cldn3** | 1.325 ± 0.332 | 2.249 ± 1.165 | 1.325 ± 0.332 | 2.249 ± 1.165 |

## 4. Discussion

Stress during the postnatal period is an important risk factor in the development of postpartum mental health disorders (Reid & Taylor, 2015; Yim et al., 2015), yet robust mouse models of postpartum depression or anxiety using postnatal stressors remain surprisingly scarce (Mundorf et al., 2022) and the potential contribution of intestinal and brain inflammatory and barrier disruptions in these models is understudied, despite evidence showing their involvement in behavioural disturbances following chronic stressors in non-postpartum contexts (Audet et al., 2011; Doney et al., 2023; Ménard et al., 2017; Russo et al., 2023). The present study examined, in C57BL/6N mice, whether MS experienced during the early postnatal period impacted maternal caregiving as well as anxiety- and depressive-like behaviours and if these were accompanied by differences in the expression of pro-inflammatory cytokines and tight junction proteins in the mPFC and the colon. Our findings first demonstrate that MS impaired the temporal pattern of maternal care behaviours but failed to alter anxiety- and depressive-like behaviours. Then, although prefrontal and colonic markers of inflammation and barrier permeability remained unchanged by MS, increased grooming time in the splash test was associated with increased transcription of TNF-α in the mPFC, but decreased transcription of IL-6 and Ocln in this region.

In contrast with the reported general disruptions in maternal care behaviours following MS (Mundorf et al., 2022), our findings suggest that specific components of maternal caregiving may differ in their susceptibility to MS depending on the stage of the postpartum period, with stressed dams taking more time to initiate contact with their pups and spending more time in passive contact with them on P3, but less time physically moving them and their nest on P7 relative to NS dams. Our findings are in line with previous reports showing that MS rat dams took more time to retrieve their first pup in the pup retrieval test on P3 (Aguggia et al., 2013; Demarchi et al., 2023; Maghami et al., 2018), supporting the possibility that MS-induced impairments in maternal responsiveness towards pups develop early during the postpartum period. In support to our finding that MS in mice may disrupt the normal trajectory of maternal care behaviour, MS mouse dams engaged less in maternal care in the early postnatal period but more later on relative to controls (Orso et al., 2018; Own & Patel, 2013). With regard to the temporal variation in maternal care impairments from P3 to P7, it is important to consider that maternal behaviour in rodents is highly organized, with certain behaviours, such as nursing and nesting, varying with the developmental stage of the pups (Weber et al., 2008). As maternal care behaviour adapts to the developmental needs of the pups (Olazábal et al., 2013), the effects of stress on caregiving may differ depending on the postpartum stage at which the behaviour is assessed.

One important finding of our study is that anxiety- and depressive-like behaviours were not affected by MS, at least when examined shortly after weaning. As previously alluded to, most of the studies that examined behavioural outcomes in MS dams were conducted in rats, with reports that did not observe changes in anxiety- and depressive-like behaviours (Aguggia et al., 2013; Bölükbas et al., 2020; Stevenson et al., 2009) and some that did report such effects (Boccia et al., 2007; Bousalham et al., 2013; Maghami et al., 2018). When compared with the very few studies conducted in mice, our findings contrast with the increases in anxiety- (Orso et al., 2018) and depressive-like (Zhang et al., 2022) behaviours reported in MS dams. Since these two studies used Balb/c mice, a mouse strain more anxious and reactive to stress than C57BL/6 mice, our findings suggest that C57BL/6 dams may be less sensitive to the behavioural consequences of MS or that these effects may not persist beyond the early postpartum period. Although there is limited evidence on the timing of mental health symptoms during the postpartum period, women appear to be most susceptible to developing depressive and anxiety disorders during the first 6 weeks after birth (Kettunen et al., 2014), with symptom onset in the first 8 weeks being associated with more severe depressive symptoms (Putnam et al., 2017), suggesting that mental health symptoms may be most pronounced during the early postnatal period. It is also possible that MS alone does not result in lasting behavioural alterations in the C57BL/6 mouse strain we used but instead increases vulnerability to subsequent stressor exposure. A study conducted by Wu et al. (2021) in C57BL/6J dams did not observe depressive- and anxiety-like behaviours at weaning with MS alone, but upon a lipopolysaccharide injection as a second “hit”, elevations in depressive- and anxiety-like behaviour were apparent, potentially pointing towards enhanced stress reactivity in the dams rather than lasting overt behavioural effects.

Pro-inflammatory cytokine elevations in the cerebrospinal fluid and the plasma have been reported in women with postpartum depression (Achtyes et al., 2020; Boufidou et al., 2009; Sha et al., 2022) but whether postnatal stressors promote and/or enhance these effects remains incompletely understood. Similar to our behavioural findings, prefrontal and colonic pro-inflammatory cytokines and tight junction proteins after weaning were not affected by MS. This contrasts with a report showing cortical reductions in IL-6 and TNF-α and changes to the gut microbiota as a result of MS in Balb/c dams (Zhang et al., 2022), again suggesting that C57BL/6 dams may be less responsive to MS. Given that tissue collection occurred on P24, 10 days following the end of the stressor and three days after weaning had been completed, it cannot be excluded that such changes occurred earlier in the postpartum period but were no longer detectable at that time.

Despite the overall absence of stress-induced behavioural effects, MS dams exhibited a non-significant trend toward spending more time in grooming episodes of 3 seconds or greater, despite no change in grooming time overall, suggesting that the stressor may have exerted subtle effects on the structure of grooming. Using a chronic social defeat stressor, Denmark et al. (2010) showed that measures capturing grooming overall, such as the total duration of grooming and number of grooming bouts, were not affected by the stressor, but differences were observed in the patterning of grooming, suggesting that chronic stressors may affect certain features of grooming behaviour rather than its overall presentation. While our study did not evaluate the microstructure of grooming, which consists of a repeated chain of stereotyped movements organized in distinct phases (Kalueff et al., 2016), the trend toward spending more time in grooming episodes of 3 seconds or more in MS dams, despite no overall difference in grooming time, could similarly reflect changes in specific grooming characteristics in the context of MS.

The associations between prefrontal markers of inflammation and barrier permeability and grooming time suggest a potential role for TNF-α, IL-6, and Ocln in this brain region in modulating grooming behaviours, irrespective of stressor exposure. Transgenic mice with a central nervous system-specific overexpression of TNF-α showed increased grooming behaviour, and this was accompanied by reduced immunoreactivity of tyrosine hydroxylase (Aloe & Fiore, 1997), which may suggest the involvement of dopamine in this behavioural phenotype. This is consistent with the observation that drug-induced antagonism of TNF-α in depressed patients results in greater motivation and willingness to exert effort for a reward (Treadway et al., 2024). Given that grooming is in part regulated by the striatum and limbic circuitry involved in motivation, affective state, and motor sequencing (Kalueff et al., 2016), evidence suggesting that TNF-α is implicated in effort-based motivation may further support our finding that TNF-α is positively associated with grooming behaviour.

A limitation of our study is that the assessment of depressive- and anxiety-like behaviours was conducted shortly after weaning, approximately 10 days following the completion of the stressor, by which time the postpartum period could have concluded. This was done primarily to avoid introducing the potential confound of additional prolonged separations from the litter during behavioural testing but limited the ability to observe postpartum-specific depressive- and anxiety-like effects. As such, it cannot be excluded that changes in these behaviours may have been present throughout the early postpartum period but had resolved by the time behavioural testing was conducted. Likewise, tissue collection occurred on P24, and it may also be possible that pro-inflammatory cytokine and tight junction protein transcription was no longer affected beyond the early postpartum period. Furthermore, maternal care behaviours were evaluated only at P3 and P7 following a 5-minute reunion, thereby limiting the assessment of dynamic changes in maternal caregiving throughout the postpartum period. While this allows observation of acute reunion behaviours, it may not be representative of broader disruptions in maternal care behaviour observed in other studies using longer recordings (Rombaut et al., 2023).

## 5. Conclusion

Overall, MS had selective effects on parameters of maternal caregiving that vary by the postnatal day assessed, suggesting dynamic changes in disruptions to maternal care behaviours in the early postpartum period. Changes in anxiety- and depressive-like behaviours and in pro-inflammatory cytokine and tight junction protein expression in the mPFC and colon in MS dams were absent, which may indicate that C57BL/6N dams may lack sensitivity to this specific MS protocol and/or that earlier, transient effects were not observed due to the use of a single assessment timepoint. Future studies using mice should examine the temporal progression of maternal care behaviours throughout the postpartum period and employ multiple cohorts to assess whether there are transient changes in affective behaviours and in gut-brain markers of inflammation and barrier permeability.

## Author contributions

VM and MCA designed the experiment. VM and TT conducted the experiment and performed the behavioural, molecular (RT-qPCR), and statistical analyses. VM and MCA interpreted the data and wrote the manuscript, which was edited and approved by all authors before submission.

## Acknowledgments

This work was supported by the Natural Sciences and Engineering Research Council (NSERC #RGPIN-2016-06146 Discovery Grant to MCA). The Canadian Institutes for Health Research (Canada Graduate Scholarship – Master’s Program to VM), NSERC (Undergraduate Student Research Award to TT), and the Parviz Sabour MSc Scholarship in Nutrition and Mental Health (MSc Scholarship to VM) also supported this research. We would like to thank the Animal Care and Veterinary Service and the Animal Behaviour and Physiology Core at the University of Ottawa.

---

**Supplementary Table 1.**
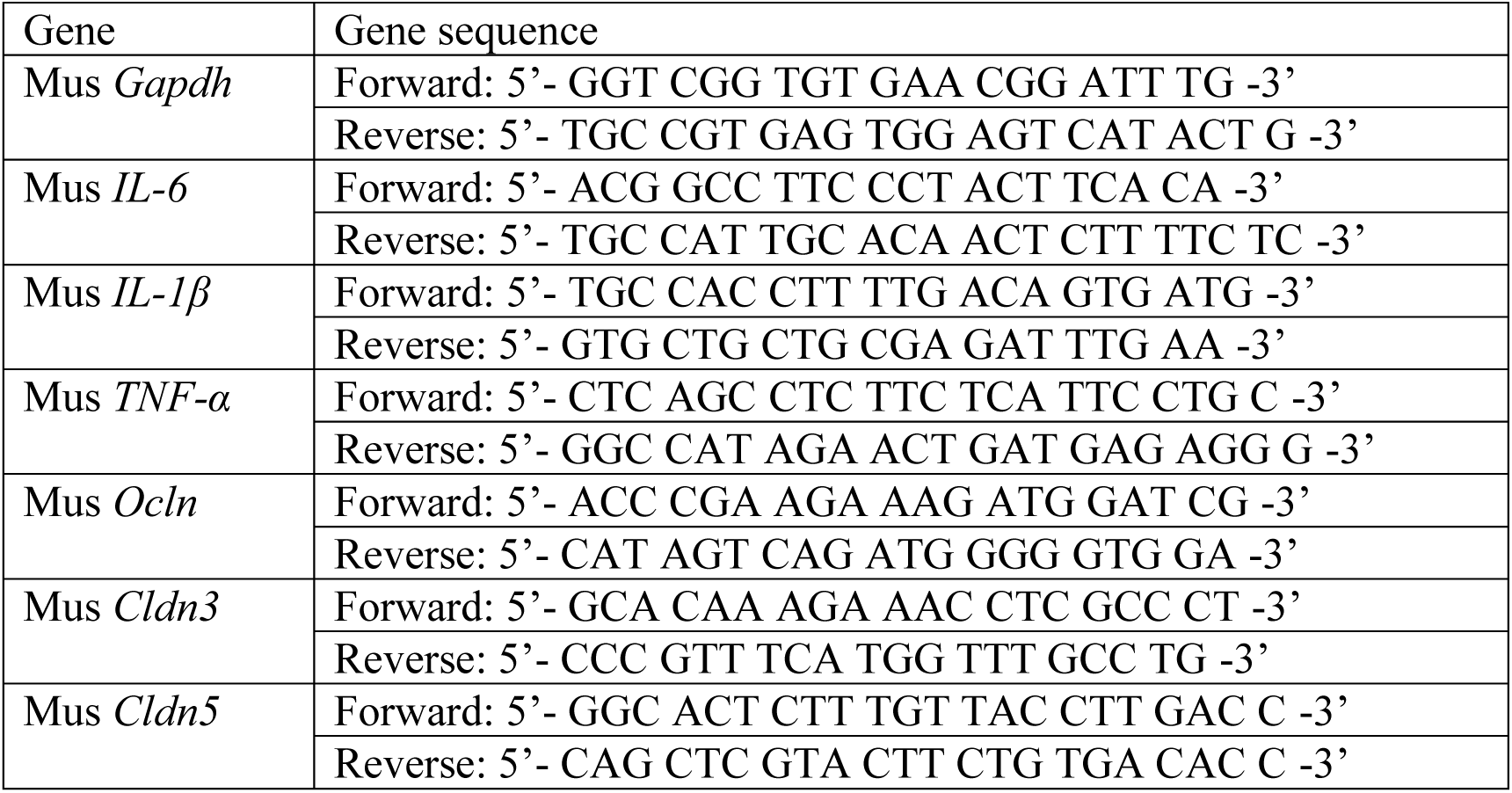
Primer sequences used in RT-qPCR analyses.

